# Modeling Decision-Making Under Uncertainty with Qualitative Outcomes

**DOI:** 10.1101/2024.08.27.609831

**Authors:** Nachshon Korem, Duek Or, Jia Rounan, Emily Wertheimer, Sierra Metviner, Michael Grubb, Ifat Levy

## Abstract

Modeling decision-making under uncertainty typically relies on quantitative outcomes. Many decisions, however, are qualitative in nature, posing problems for traditional models. Here, we aimed to model uncertainty attitudes in decisions with qualitative outcomes. Participants made choices between certain outcomes and the chance for more favorable outcomes in quantitative (monetary) and qualitative (medical) modalities. Using computational modeling, we estimated the values participants assigned to qualitative outcomes and compared uncertainty attitudes across domains. Our model provided a good fit for the data, including quantitative estimates for qualitative outcomes. The model outperformed a utility function in quantitative decisions. Additionally, we found an association between ambiguity attitudes across domains. Results were replicated in an independent sample. We demonstrate the ability to extract quantitative measures from qualitative outcomes, leading to better estimation of subjective values. This allows for the characterization of individual behavior traits under a wide range of conditions.

**Author Summary:** In the current study, we explored how people make decisions when the outcomes aren’t easily measured in numbers, such as in medical choices. Traditional mathematical models, which rely on numerical data, often fall short in these situations, leading to a gap in understanding how people evaluate these qualitative outcomes. Using hierarchical Bayesian modeling, we developed a model that bridges this gap by translating qualitative outcomes into individualized quantitative values, enabling us to better understand the underlying decision-making processes. Our model not only provides a better fit to real-world data than existing models with qualitative or quantitative outcomes but also allows for meaningful comparisons of how people handle uncertainty across different decision-making scenarios. This approach opens new doors for studying decision-making in areas where traditional methods struggle, offering a more nuanced view of human behavior in complex situations.

## Introduction

Life is a series of decisions where most outcomes are uncertain. These decisions range from trying a new dish at our favorite restaurant to selecting a life-saving medical treatment. Often, these decisions are quantitative in nature; for example, buying a lottery ticket for $2 with a 1 in 300 million chance of winning $21 million. Many decisions, however, are qualitative, such as choosing whether to dine at an Italian or Thai restaurant. The ability to compare different qualitative outcomes implies that we can derive some form of comparable subjective value for these outcomes [1]. Here, we aimed to quantify how qualitative outcomes affect uncertainty attitudes.

Uncertainty around choice outcomes can be categorized into two components: risk and ambiguity. Risk occurs when probabilities of potential outcomes are precisely known [2]; ambiguity refers to situations where these probabilities are partially or entirely unknown [3]. Prior research has shown that individuals generally exhibit an aversion to both risk and ambiguity in scenarios involving potential gains [4–7], but that these attitudes vary substantially across individuals and are not strongly correlated with each other [7–13,13]. Many studies characterized these attitudes using choices with monetary (or point) outcomes [5,8,10,12,14–16]. The few studies that quantified individual attitudes in other domains still employed quantitative outcomes, including numbers of M&Ms and milliliters of water [1], months of extended lifespan [17], or milligrams of medication, and minutes spent with social partners [18]. Although some earlier studies did examine qualitative decision-making [19,20], there is still no straightforward way to model these kinds of decisions. As a result, such decisions are typically not addressed in neuroimaging and psychiatric research.

In this study, we estimated risk and ambiguity attitudes in two separate modalities: quantitative (monetary decisions) and qualitative (medical decisions), leveraging computational modeling to extract values from qualitative outcomes and examine how uncertainty attitudes influence decision-making across various domains. Sixty-six in-person and 332 online participants engaged in a task of decision-making under uncertainty, where they made choices between a certain outcome and the chance for a more favorable outcome. Our objectives were to (1) estimate subjective values for a range of qualitative outcomes, (2) assess the model’s fit using quantitative outcomes for comparison, and (3) explore how attitudes towards uncertainty vary across different domains.

## Method

### Study 1 (in-person)

#### Participants

One hundred and one adults (48 females; age range = 18–89; mean 52.97, SD ±22.41) were screened for the experiment. All participants were screened over the phone to ensure the absence of major medical conditions, including neurological illness and lifetime Axis I psychiatric disorders. Participants provided written consent after a detailed explanation of the study was provided, following the institute guidelines approved by the Yale Human Investigation Committee. Ten participants did not complete the study and were not included in the analysis, resulting in a sample of ninety-one participants (failed to complete the task n=4; failed to come to consecutive sessions n=6).

To ensure all participants were cognitively healthy, we administered the Montreal Cognitive Assessment (MoCA) [21], which can detect mild cognitive impairments. Data from participants who scored less than 26 on the test were excluded [22], resulting in a sample of seventy-one cognitively healthy participants (32 females; age range = 18–88; mean 49.68 ±22.3 SD).

#### Procedure

Participants in this study came for three sessions completed within one week. Task data reported here were collected on the first session. The MoCA was completed in session 3, along with several other questionnaires. In brief, in session 1, participants completed the decision-making task and a reversal reinforcement learning task that is not reported here. In session 2, participants completed an fMRI task. Finally, in session 3, participants completed several questionnaires assessing cognitive ability and general IQ. Participants were paid for each session separately and an extra fee for successfully completing the experiment.

##### Decision-making task

###### Risk and ambiguity in the monetary domain

The task was based on a previously developed task [12] used in multiple studies. On each trial, participants chose between a small certain gain ($5) and a lottery that offered a larger amount. The lottery was risky in half of the trials, i.e., with known outcome probability. The risky lotteries were represented as bags containing red and blue chips. The numbers of red and blue chips were indicated by the percentage of a rectangle colored in red and blue and the numbers on the bag. Three different outcome probabilities were used (25%, 50%, and 75%). Dollar amounts ($5, $8, $12, and $25) next to each color (Figure 1a) indicated the amount of money that could be won if a chip of that color was drawn. In the remaining trials, the lottery was ambiguous, i.e., outcome probability was not precisely known. Ambiguity was achieved by occluding part of the bag (Figure 1b), rendering the probability of drawing a chip of a certain color partially unknown. Increasing the occluder size (24%, 50%, and 74%) increases the ambiguity level or the range of possible probabilities for drawing a red or blue chip. Each combination (amount, risk/ambiguity) was repeated four times. On twelve (out of 84) trials, participants were asked to choose between $5 for sure and a chance to win $5. Those trials were used as attention checks. Participants who failed six or more attention checks (n=3) were removed from the analysis in the risk and ambiguity task. In addition, participants who chose the lottery less than two times were omitted from the analysis because their data could not be fitted with a model (n=2). This exclusion was necessary because our choice function requires response variability; a consistent choice of one option provides no data points for model estimation. The final sample included 66 participants. At the end of the experiment, the computer randomly selected one of the trials, and the participants acted out the trials by selecting a chip. However, they did not receive the additional payment. We opted for a hypothetical outcome to make the monetary and medical conditions similar to each other (see below).

**Figure 1.**
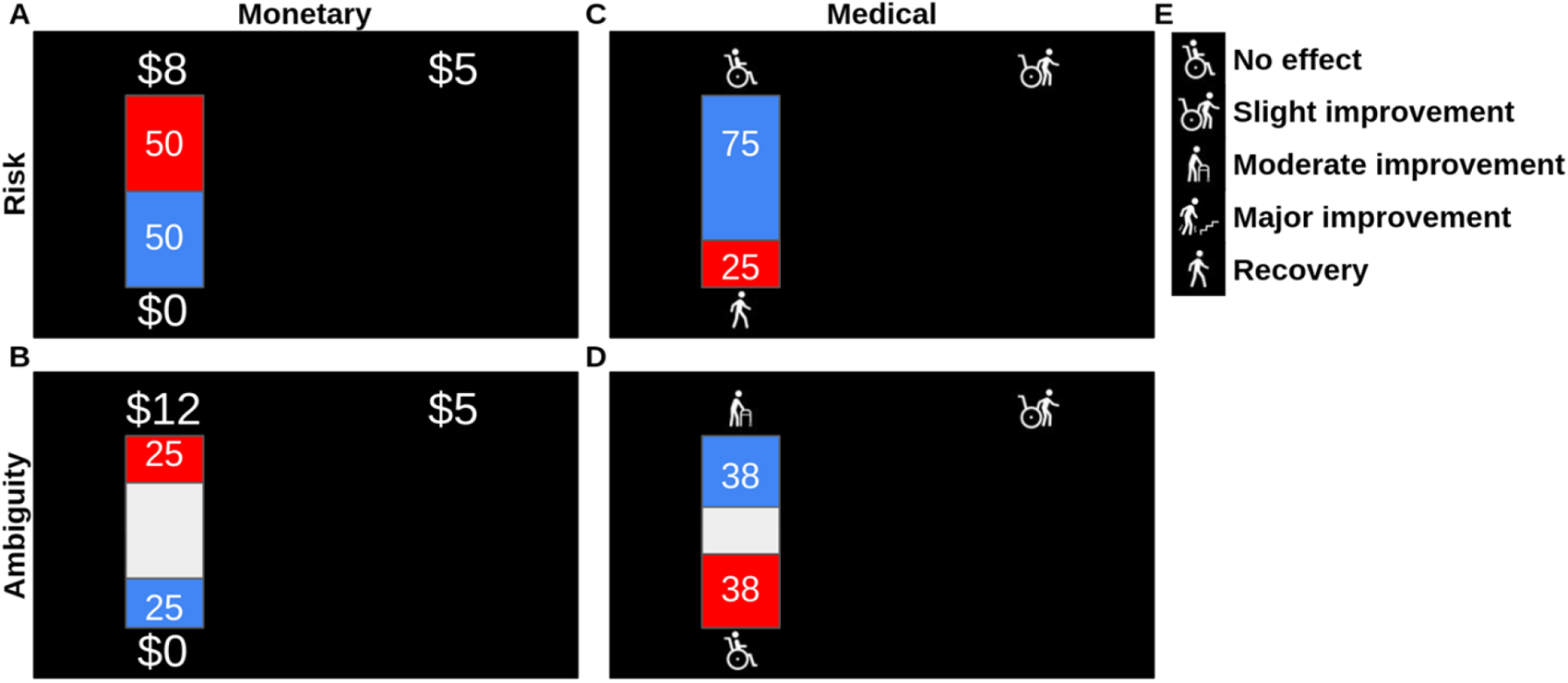
Risk and ambiguity task Task Design: Participants were presented with choices between an uncertain option and a certain outcome across four scenarios: risky monetary decisions (A), ambiguous monetary decisions (B), risky medical decisions (C), and ambiguous medical decisions (D). In the risky scenarios (A, C), the outcome probabilities were visually represented by red and blue rectangles, and these probabilities were fully disclosed to the participants. In the ambiguous scenarios (B, D), the probability information was partially obscured by a grey rectangle, indicating uncertainty. The outcome probabilities in risky trials were set at 25%, 50%, and 75%, while the levels of ambiguity (indicated by the grey area) were set at 74%, 50%, and 24%. There were four possible outcomes for monetary decisions ($5, $8, $12, and $25) and four potential medical outcomes (E). Each unique pairing of uncertainty and outcome levels was presented to the participants four times.

###### Risk and ambiguity in the medical domain

Participants were presented with a hypothetical scenario in which they were involved in a car accident and, as a result, suffered a spinal injury (supplementary methods for more details). They were asked to choose between two medical treatments (Figure 1c, 1d), a known treatment with a known outcome (“slight improvement,” parallel to a fixed monetary gain of $5) or the experimental treatment (Figure 1e), with outcomes varying in the level of improvement (“moderate,” “major,” or “complete recovery”). The likelihood of the outcome of the experimental treatment varied with different levels of risk and ambiguity (parallel to playing a lottery). Outcome probabilities and ambiguity levels were the same as in the monetary task and were presented graphically and verbally. All aspects of the experimental design were similar to the monetary task. The order of the monetary and medical tasks was counterbalanced across participants.

### Study 2 (online)

This dataset was used to confirm and replicate the results. This is a secondary analysis of a previously published dataset [23].

#### Participants

Four hundred and four adults (212 females; age range = 20–80; mean 49.362, SD ±14.849) were recruited using Amazon Mechanical Turk (mTurk). Participants provided consent online after reading a detailed explanation of the study following Yale Human Investigation Committee guidelines. Seventy-two participants did not complete the study and were not included in the analysis, resulting in a sample of three-hundred and thirty-two participants.

#### Procedure

Participants completed tasks similar to those described above, with two exceptions. First, each lottery was presented twice instead of four times. Second, an additional condition of 100% ambiguity was introduced. For more details, please see Xu et al., [23].

### Modeling approach

We employed hierarchical Bayesian modeling to capture and examine subjects’ choice behavior. By leveraging hierarchical Bayesian modeling (HBM), we were able to uncover hidden variables that offer valuable insights into the underlying mechanisms of decision-making. HBMs allow for partial pooling of data across the population, meaning that individual data points contribute to both individual and group-level estimates. This results in more robust and accurate posterior distributions compared to non-hierarchical models, particularly when dealing with small sample sizes or individual-level variability [24]. Moreover, we were able to compare different models to assess which described the data better. In the context of monetary decisions, to model the subjective value assigned to each option, we utilized a power (utility) function [6] with a linear effect of ambiguity on the perceived probability (Equation 1 – Classic Utility Model) [12].

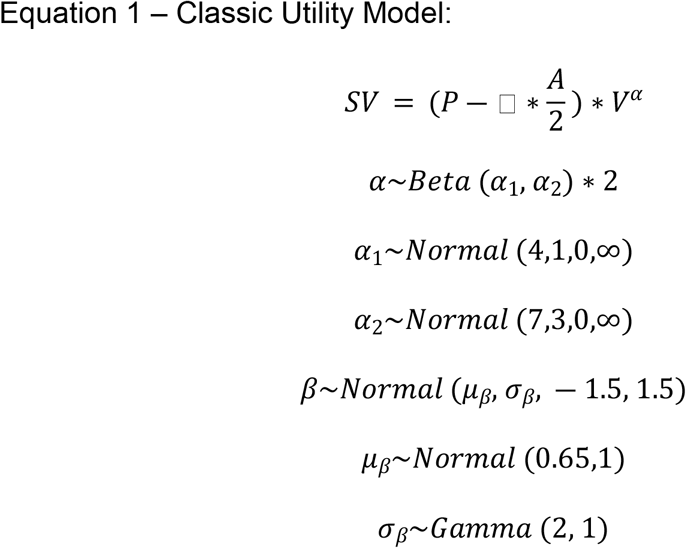

We modeled the subjective value (SV) using the objective probability (P; for risky options P = 0.25, 0.5, or 0.75, for ambiguous options P = 0.5, and for the certain gain P = 1), ambiguity level (A; for risky or certain options A= 0, for ambiguous trials A = 0.24, 0.5, or 0.74), and potential winnings (V; 5, 8, 12, 25 dollars for the lotteries and 5 dollars for the sure bet). Risk attitude (α) and ambiguity attitude (β) were incorporated to capture individual differences in risk and ambiguity preferences. Hyperpriors are higher-level distributions that set prior beliefs on the parameters of the model’s primary prior distributions. They are beneficial as they allow for incorporating prior knowledge and uncertainty about the parameters, leading to more flexible and robust models. Importantly, the same priors were used for the two independent data sets. Hyperpriors were chosen based on previous data [12], with slight risk (mean 0.72) and ambiguity (mean 0.65) aversion (see supplementary results for comparison with less and uninformed hyper-priors).

To estimate the probability of choosing the lottery on each trial, we fitted a logistic choice function (equation 2), in which *γ* is the inverse temperature parameter.

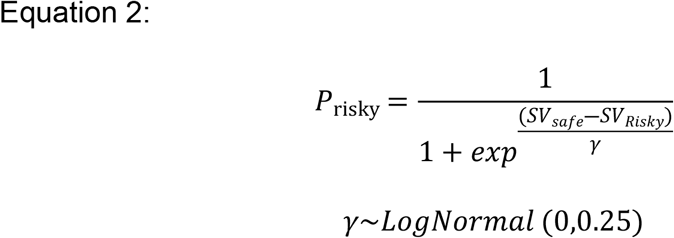

In addition, we tested a “trembling-hand” logistic choice function (Equation 3), where there is less dependency between the risk and ambiguity parameters and the slope of the logistic function [25].

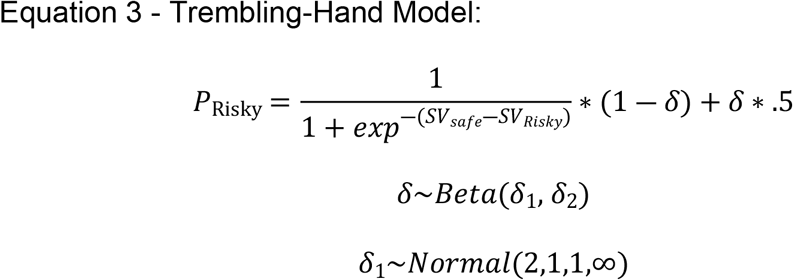

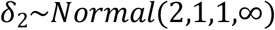

In the medical domain, fitting the utility function is challenging due to the qualitative nature of outcomes, which lacked a quantifiable value for the value (V) component of the model. To tackle this issue, we used the model to estimate the subjective value associated with each outcome. Notably, we excluded the risk aversion parameter, as subjective values were individually tailored to each outcome and participant (equation 4 – Estimated Value Model).

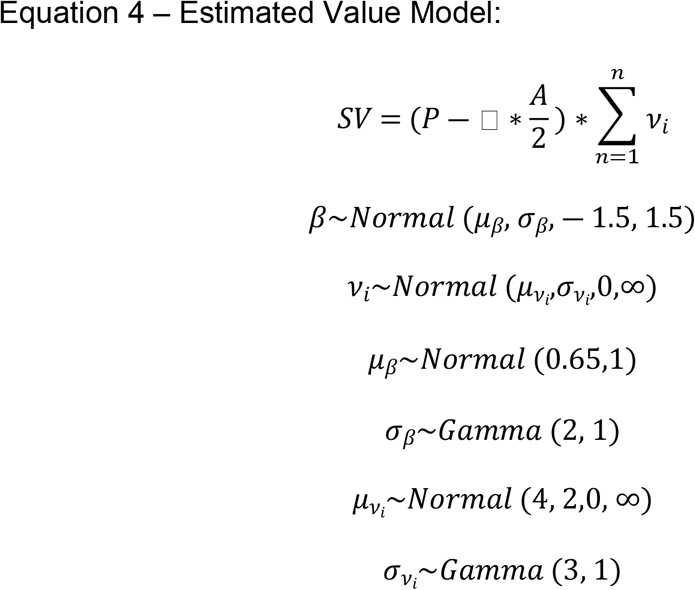

In this model, we estimated the subjective value (*ν*) of each outcome (i) based on the cumulative values of the preceding outcomes. We assumed the outcomes were ordinal (e.g., slight improvement < moderate improvement), and thus, we modeled the subjective value of each outcome as the sum of the subjective value of the preceding outcomes plus an additive value representing the improvement.

To evaluate the model’s fit, we introduced a baseline model devoid of subject-specific parameters, serving as a ‘straw man’ to establish a reference point for the estimated model’s performance. In the medical domain, we utilized the category level (e.g., 1 for slight improvement, 2 for moderate improvement) as the value (V) in the model (equation 5).

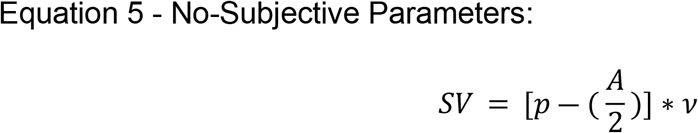

All models converge with rHat<1.01 and effective sampling rate > 1000. All analyses were conducted in Python 3.10.14, utilizing the ‘PyMC’ (version 4.1.7) [26] and ‘ArviZ’ (version 0.17.1) [27] packages. We utilized the No-U-Turn Sampler (NUTS) for Markov chain Monte Carlo (MCMC) inference, adhering to PyMC’s default settings: 1000 draws, 1000 tuning steps, no thinning, and an 80% acceptance rate. The code can be found here: https://github.com/KoremNSN/QualMod/tree/main

#### Model comparison

The different models were compared using a <u>Leave-One-Out</u> (LOO) cross-validation method to estimate out-of-sample predictive fit [28]. This method partitions the data into train and test data and fits the training-based data on the holdout test data to evaluate the fit. This process is done repeatedly. We used the ‘Arviz’ implementation to compute the LOO of the models. Unlike Log Likelihood, LOO measures expected log pointwise predictive density (ELPD). Thus, higher LOO points to a better fit.

#### Simulations

The estimated model has more degrees of freedom. Hence, a better fit could result from overfitting [29]. One way to ensure that the results represent real phenomena rather than overfit is by using simulations [30]. Using the Utility function (Equation 1) and the Estimated Value (Equation 4) models, we tested several scenarios, including different numbers of participants and levels of noise. The simulation assumed that the participant uses the utility function to make choices. However, each choice introduces a unique noise level to the risk attitude (α) and ambiguity attitude (β) parameters. The noise came from a normal distribution with a mean of 0 and a standard deviation that changed based on the simulation. To ensure model convergence, the risk attitude was limited to a range between 0.1 and 1.6, and the ambiguity attitude to a range between -1.4 and 1.4. We tested the following sets of parameters (N, noise): (30, 0.1), (30, 0.3), (30, 0.5), (60, 0.3), (60, 0.5), (120, 0.5), (300, 0.5). LOO values and weight were used to compare model fit.

## Results

Sixty-six participants (Table 1 for demographics) completed a computerized decision-making task under uncertainty with qualitative and quantitative outcomes. Our objectives were to determine if we could (1) extract subjective values from choices between options with qualitative outcomes, (2) assess how well these estimated values fit the observed data, and (3) explore associations across uncertainty attitudes. Data from an additional sample of 332 online participants was used to replicate the results.

**Table 1.** Demographic description of the sample.

|  | In-person sample<br>N / Mean (SD); min-max | Online sample<br>N / Mean (SD); min-max |
| --- | --- | --- |
| Age | 49.14 (22.76); 18-88 | 49.49 (14.73); 20-80 |
| Sex (females) | 30 (45.45%) | 164 (49.40%) |
| MoCA | 28.39 (1.28); 26-30 |  |
| Education |  |  |
| High school graduate, GED, or less | 4 | 27 |
| Some college or post-high school | 1 | 85 |
| College Graduate | 22 | 139 |
| Advanced graduate or professional degree | 27 | 44 |
| Estimated household income |  |  |
| \$14,999 or less | 12 | 26 |
| \$15,000-24,999 | 3 | 21 |
| \$35,000-49,999 | 11 | 103 |
| \$50,000-74,999 | 8 | 93 |
| \$75,000-99,999 | 4 | 37 |
| \$100,000-149,999 | 13 | 37 |
| 150,000-250,000 | 11 | 11 |
| \$250,000-350,000 | 2 | 2 |
| \$350,000 or more | 1 | |
| missing | 1 | 2 |
| History of major surgery |  |  |
| Yes | 29 |  |
| No | 36 |  |
| missing | 1 |  |

### Extracting Quantitative Values from Qualitative Outcomes

We applied the Estimated Value model (equation 4) to the medical decision data to obtain estimated subjective values for the different levels of qualitative outcomes. For comparison, we also applied the No-Subjective Parameters model (equation 5) to the data (see Table 2 for model comparison). The Estimated Value model demonstrated a superior fit to the data in both samples. Subsequently, we extracted the mean added value for each category assigned by each participant. This approach provided estimates of the added values people attribute to the various outcomes. The mean value for slight improvement was 6.93, with a standard deviation (SD) of 1.79. This value increased by 9.00 (SD 2.62) for moderate improvement, by an additional 7.04 (SD 2.89) for major improvement, and finally by 4.18 (SD 2.41) for complete recovery (Figure 2A). For the online sample, the mean value for slight improvement was 8.63 (SD 2.54), which increased by 12.62 (SD 4.25) for moderate improvement, by an additional 4.66 (SD 2.56) for significant improvement, and finally by an additional 2.37 (SD 1.61) for complete recovery.

**Table 2.**
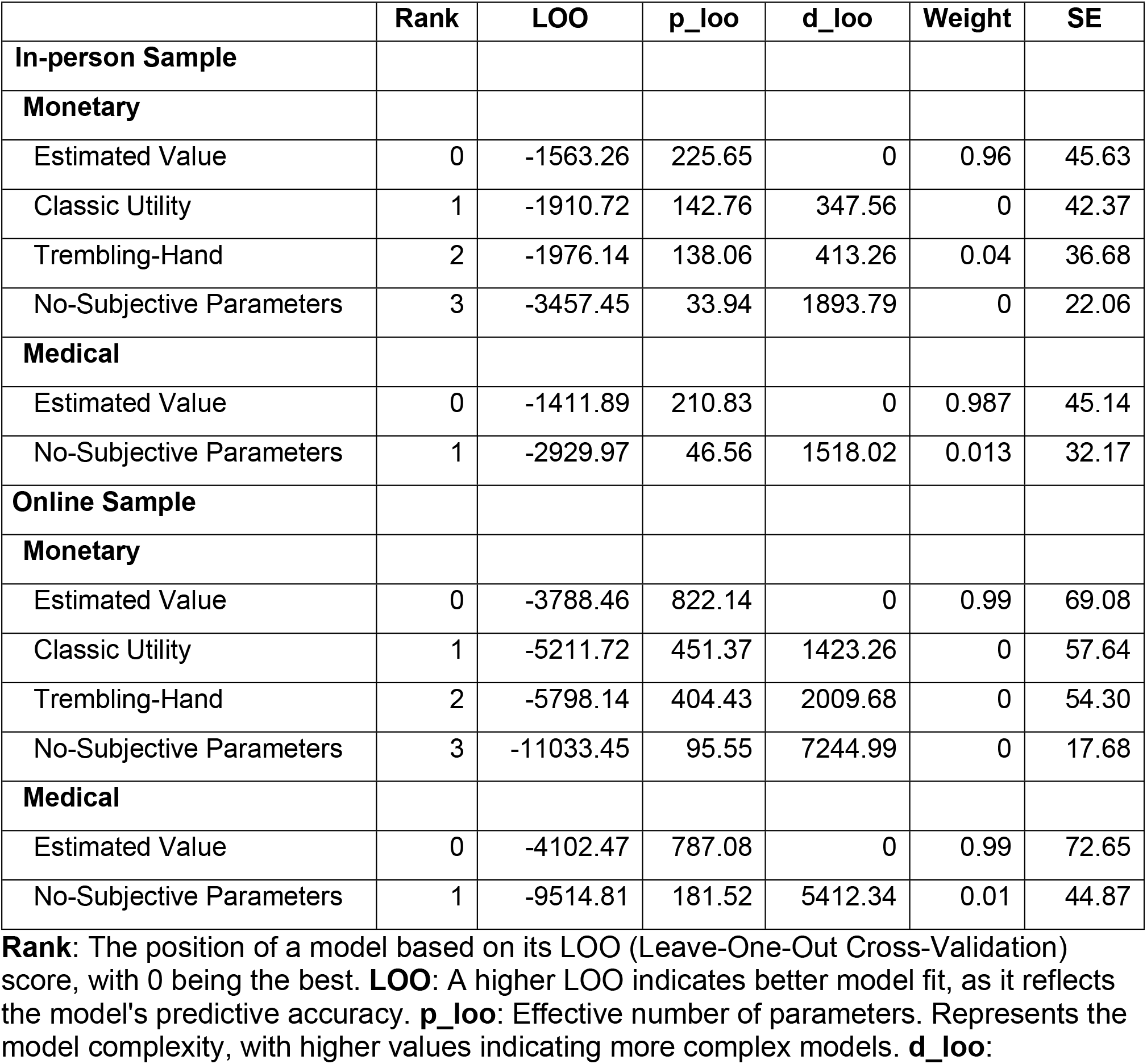
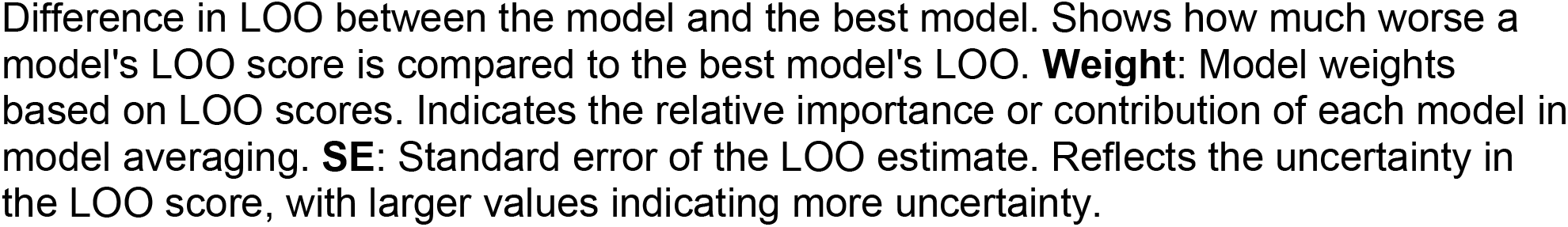
Model comparison.

**Figure 2.**
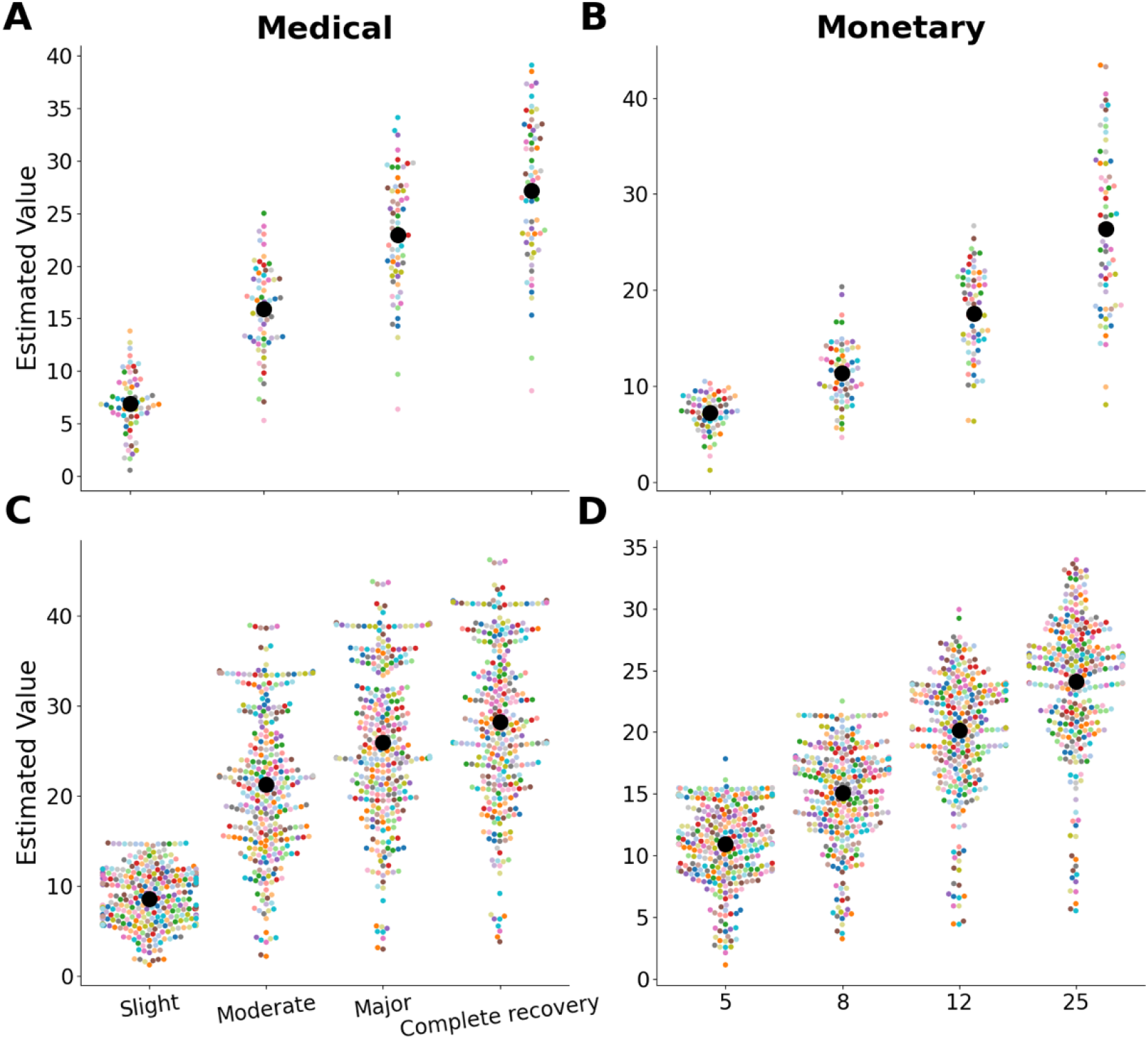
Estimated value compared with actual values and categories. The estimated values from the Estimated Value model for each category. **Panes A and B** show the estimated values for the medical decision-making task (Pane A) and the monetary decision-making task (Pane B) for the in-person sample. **Panes C and D** show the estimated values in the medical decision-making task (Pane C) and the monetary decision-making task (Pane D) for the online sample. Each colored dot represents an individual participant, while the large black dot indicates the mean estimated value for each category.

### Model Validation and Comparative Analysis

We applied the Estimated Value model to the monetary decision data to facilitate a direct comparison with a classical utility model (equation 1). Using the monetary decision data, we evaluated the effectiveness of the Estimated Value model (equation 4) by comparing it to various models that incorporate an objective estimate of the outcome value. Specifically, we compared the Estimated Value model against several alternatives: a No-Subjective Parameters model (equation 5), a Classical Utility model (equation 1), and a Classical Utility with a Trembling-Hand choice function model (equation 3). Our analysis revealed that the Estimated Value model outperformed the other models in terms of fit to the data within the monetary domain in both samples (see Table 2). This finding underscores the robustness of the Estimated Value model across different decision-making contexts.

We also derived values for the monetary categories. The mean value for $5 was 7.22 (SD 1.60). This value increased by 4.13 (SD 1.47) for $8, by an additional 6.23 (SD 2.18) for $12, and finally by 8.77 (SD 3.41) for $25 (Figure 2B). For the online sample, the mean value for $5 was 10.91 (SD 2.72), which increased by 4.13 (SD 1.84) for $8, by an additional 5.04 (SD 2.35) for $12, and finally by an additional 3.92 (SD 2.18) for $25.

### Cross-domain association between uncertainty attitudes

Finally, we explored how attitudes toward uncertainty vary across different domains. Using robust regression [31,32], we found that ambiguity aversion (β) in the medical domain was strongly and positively associated with the same attitude in the monetary domain (mean slope = 0.76, 89% HDP [0.63, 0.91]; Figure 3). This association was replicated in the online sample (mean slope = 0.37, 89% HDP [0.29, 0.44]). This finding suggests that attitudes towards ambiguity are consistent across different domains. It is important to note that risk attitudes, as operationalized in our models, depend on the outcome levels; that is, the value incorporates the risk preference. Thus, it precludes a straightforward comparison of risk attitudes across domains.

**Figure 3.**
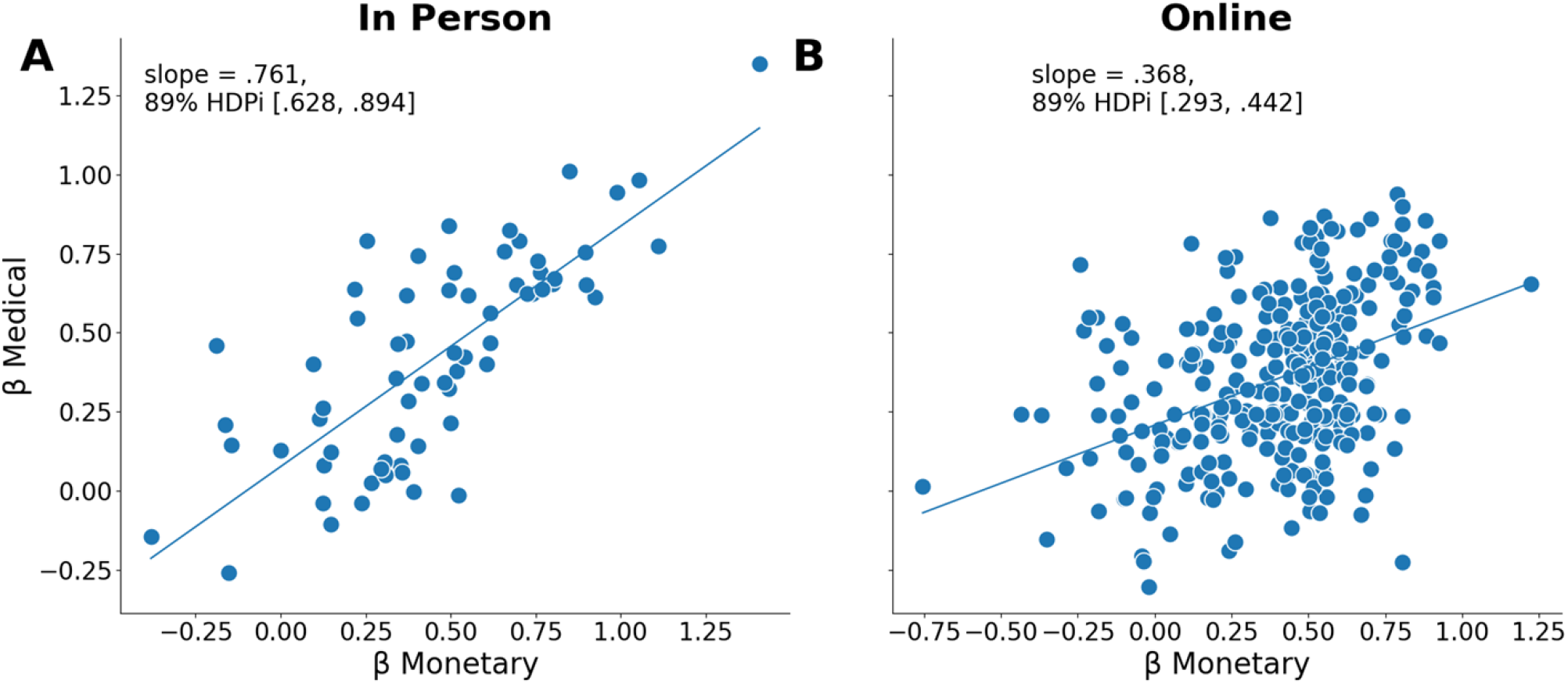
Cross-domain association in ambiguity aversion (β) The positive association between ambiguity aversion (β) in the monetary (x-axis) and medical (y-axis) domains in the in-person (**Pane A**) and online (**Pane B**) samples. The mean slope of the robust regression and the 89% highest density posterior interval (HDPi) are indicated in the panels.

#### Simulations

The simulation results demonstrate that model performance is influenced by noise levels. With lower noise (0.1), the classic utility model provided a better fit. However, as noise increased to 0.3 and 0.5, the estimated value model consistently showed superior performance, regardless of the number of subjects. Specifically, at higher noise levels, the estimated value model had higher LOO values and weights across different sample sizes. For detailed results, see supplementary results.

## Discussion

This study aimed to quantify the values of qualitative outcomes and to characterize individual uncertainty attitudes when making choices between such outcomes. Participants made a series of choices between certain outcomes and uncertain, potentially better, outcomes. Through computational modeling, we estimated the values participants assigned to different qualitative outcomes and assessed their attitudes towards ambiguity, which were consistent across two domains. Notably, our model outperformed a classical utility model with access to the objective amounts in the monetary domain. Overall, our model demonstrates a good fit and allows for better estimation of subjective values for both qualitative and quantitative outcomes.

When testing uncertainty attitudes, one challenge that often arises is how to treat qualitatively different outcomes [33]. Theoretically, this problem stems from the difficulty in comparing outcomes that do not share a common metric [34]. Methodologically, it complicates the design of experiments and the interpretation of results, as traditional approaches often rely on quantitative measures [35]. Statistically, it poses challenges in modeling and analyzing data due to the lack of a consistent scale for comparison. To address this issue, researchers often quantify an aspect of the outcome (e.g., milligrams of medication) [17,18,36] or treat the outcomes as categories with a fixed distance between them [23]. In this study, we utilized computational modeling to extract estimated values for each outcome, providing insights into the processes underlying participants’ decision-making. This approach allows for the use of different categories without the assumptions of specific relationships among them. By estimating these values, we can examine individual differences and use them in more precise parametric analyses of brain responses to value, enhancing our understanding of the neural mechanisms involved in processing uncertainty.

Uncertainty is especially relevant in the medical domain, as it is central to health decisions across the entire continuum of medical care [37,38]. While medical diagnoses and treatments are often described qualitatively, experiments assessing uncertainty attitudes typically quantify these outcomes into measures like years of life [17,39] or milligrams of medication [18]. This simplification abandons the original complex qualitative outcome. In contrast, our model allows for the introduction of complete qualitative outcomes. Our focus on ambiguity - rather than risk - attitudes in this study warrants an explanation. Technically, in the Estimated Value model, risk attitudes are not quantified separately but are integrated into the estimated values (Equation 4). More broadly, any risk attitude measure relies on the outcomes and requires equating the outcomes across domains. In contrast, ambiguity attitudes measure how ambiguity affects choices within a domain, and this modulation can be compared across domains. Previous studies suggest that subtle experimental manipulations can alter decision attitudes [40] and that ambiguity attitudes are less stable over time [11,23] and lack a known structural correlate [41,42]. Nevertheless, our results indicate that ambiguity attitudes are consistent across domains when studied simultaneously. Future studies should explore disentangling the risk component from the estimated values.

In the monetary domain, we demonstrated that our Estimated Value model fits the data better than a classical utility model. While the utility model we used is constrained to follow a specific curve [43], our estimated model has more degrees of freedom and can adapt more flexibly to the data. To mitigate the risk of overfitting, we employed a leave-one-out cross-validation procedure to compare the models. Our simulations also support our modeling approach. We simulated hypothetical participants who made choices based on the Classical Utility Model. As expected, in scenarios with low noise levels, the Classical Utility model outperformed the Estimated Value model. This result highlights how the LOO algorithm effectively penalizes the Estimated Value model for its extra complexity. However, as noise levels increased, leading to a higher deviation from the utility function, the Estimated Value model consistently outperformed the Classical Utility model, regardless of the number of participants. This suggests that the Estimated Value model is more adaptable to noisy data, providing a better fit for real participants’ behavior.

The estimated model may be better equipped to capture phenomena such as the framing effect or range-frequency theory [44–46]. Unlike the utility function, which assumes outcomes are sampled from a single curve describing a person’s behavior, the estimated model allows for unique curvatures for each set of outcomes. This means that the lowest and highest amounts create a frame of reference against which all other outcomes are compared. By learning about the possibilities within this frame, participants adjust their expectations accordingly. This flexibility enables the estimated model to more accurately reflect how people perceive and evaluate different outcomes in varying contexts.

Having both monetary (quantitative) and medical (qualitative) datasets allowed us to assess the Estimated Value model with quantitative data and gain confidence in the model’s ability to extract values in the qualitative dataset. The model’s fit on quantitative data provides evidence that the estimates for the qualitative outcomes represent participants’ true values. While these estimates are on a relative scale, without specific measurement units, they open the possibility for use in future studies to examine value representation in the brain. Additionally, these estimates can be applied to evaluate the values of outcomes across different scenarios, enhancing our understanding of how people perceive and compare various types of outcomes.

To conclude, we present a model capable of assigning quantitative values to qualitative outcomes. The model demonstrates a better fit for both qualitative and quantitative data compared to other potential models. Although more complex than a classical utility function, both model comparisons and simulations suggest that the improved fit is not due to overfitting. This model opens new avenues for exploring the relationships between different domains, outcomes that cannot be objectively quantified, and the representation of value in the brain.

## Notes

### Competing Interest Statement

The authors have declared no competing interest.

## References

1. Levy DJ, Glimcher PW. The root of all value: a neural common currency for choice. Curr Opin Neurobiol. 2012;22: 1027–1038. doi:10.1016/j.conb.2012.06.001

2. Glimcher PW. Understanding risk: A guide for the perplexed. Cogn Affect Behav Neurosci. 2008;8: 348–354. doi:10.3758/CABN.8.4.348

3. Ellsberg D. Risk, Ambiguity, and the Savage Axioms. Q J Econ. 1961;75: 643. doi:10.2307/1884324

4. Camerer C, Weber M. Recent developments in modeling preferences: Uncertainty and ambiguity. J Risk Uncertain. 1992;5: 325–370.

5. Hsu M, Bhatt M, Adolphs R, Tranel D, Camerer CF. Neural Systems Responding to Degrees of Uncertainty in Human Decision-Making. Science. 2005;310: 1680–1683. doi:10.1126/science.1115327

6. Kahneman D, Tversky A. Prospect Theory: An Analysis of Decision under Risk. Econometrica. 1979;47: 263–291. doi:10.2307/1914185

7. Tymula A, Rosenberg Belmaker LA, Ruderman L, Glimcher PW, Levy I. Like cognitive function, decision making across the life span shows profound age-related changes. Proc Natl Acad Sci. 2013;110: 17143–17148.

8. Cohen M, Jaffray J-Y, Said T. Experimental comparison of individual behavior under risk and under uncertainty for gains and for losses. Organ Behav Hum Decis Process. 1987;39: 1–22. doi:10.1016/0749-5978(87)90043-4

9. FeldmanHall O, Glimcher P, Baker AL, Phelps EA. Emotion and decision-making under uncertainty: Physiological arousal predicts increased gambling during ambiguity but not risk. J Exp Psychol Gen. 2016;145: 1255–1262. doi:10.1037/xge0000205

10. Huettel SA, Stowe CJ, Gordon EM, Warner BT, Platt ML. Neural signatures of economic preferences for risk and ambiguity. Neuron. 2006;49: 765–775. doi:10.1016/j.neuron.2006.01.024

11. Konova AB, Lopez-Guzman S, Urmanche A, Ross S, Louie K, Rotrosen J, et al. Computational Markers of Risky Decision-making for Identification of Temporal Windows of Vulnerability to Opioid Use in a Real-world Clinical Setting. JAMA Psychiatry. 2020;77: 368–377. doi:10.1001/jamapsychiatry.2019.4013

12. Levy I, Snell J, Nelson A, Rustichini A, Glimcher P. Neural Representation of Subjective Value Under Risk and Ambiguity. J Neurophysiol. 2010;103: 1036–1047. doi:10.1152/jn.00853.2009

13. Tobler PN, Christopoulos GI, O’Doherty JP, Dolan RJ, Schultz W. Risk-dependent reward value signal in human prefrontal cortex. Proc Natl Acad Sci. 2009;106: 7185–7190. doi:10.1073/pnas.0809599106

14. Blankenstein NE, Crone EA, van den Bos W, van Duijvenvoorde ACK. Dealing With Uncertainty: Testing Risk- and Ambiguity-Attitude Across Adolescence. Dev Neuropsychol. 2016;41: 77–92. doi:10.1080/87565641.2016.1158265

15. Peysakhovich A, Karmarkar UR. Asymmetric Effects of Favorable and Unfavorable Information on Decision Making Under Ambiguity. Manag Sci. 2016;62: 2163–2178. doi:10.1287/mnsc.2015.2233

16. Serra D. Decision-making: from neuroscience to neuroeconomics—an overview. Theory Decis. 2021;91: 1–80. doi:10.1007/s11238-021-09830-3

17. Attema AE, Bleichrodt H, L’Haridon O. Ambiguity preferences for health. Health Econ. 2018;27: 1699–1716. doi:10.1002/hec.3795

18. Seaman KL, Gorlick MA, Vekaria KM, Hsu M, Zald DH, Samanez-Larkin GR. Adult age differences in decision making across domains: Increased discounting of social and health-related rewards. Psychol Aging. 2016;31: 737.

19. Doyle J, Thomason RH. Background to Qualitative Decision Theory. AI Mag. 1999;20: 55– 55. doi:10.1609/aimag.v20i2.1456

20. Dubois D, Godo L, Prade H, Adriana Z. Making Decision in a Qualitative Setting: from Decision under Uncertaintly to Case-based Decision. 1998. p. 607.

21. Nasreddine ZS, Phillips NA, Bédirian V, Charbonneau S, Whitehead V, Collin I, et al. The Montreal Cognitive Assessment, MoCA: A Brief Screening Tool For Mild Cognitive Impairment. J Am Geriatr Soc. 2005;53: 695–699. doi:10.1111/j.1532-5415.2005.53221.x

22. Rossetti HC, Lacritz LH, Cullum CM, Weiner MF. Normative data for the Montreal Cognitive Assessment (MoCA) in a population-based sample. Neurology. 2011;77: 1272– 1275.

23. Xu CY, Dan O, Jia R, Wertheimer E, Chawla M, Fuhrmann-Alpert G, et al. Quantitative vs. Qualitative Outcomes: A Longitudinal Study of Risk and Ambiguity in Monetary and Medical Decision-Making. 2024. doi:10.21203/rs.3.rs-4249490/v1

24. Kruschke J. Doing Bayesian Data Analysis: A Tutorial with R, JAGS, and Stan. Academic Press; 2014.

25. Krefeld-Schwalb A, Pachur T, Scheibehenne B. Structural parameter interdependencies in computational models of cognition. Psychol Rev. 2022;129: 313–339. doi:10.1037/rev0000285

26. Abril-Pla O, Andreani V, Carroll C, Dong L, Fonnesbeck CJ, Kochurov M, et al. PyMC: a modern, and comprehensive probabilistic programming framework in Python. PeerJ Comput Sci. 2023;9: e1516.

27. Kumar R, Carroll C, Hartikainen A, Martín OA. ArviZ a unified library for exploratory analysis of Bayesian models in Python. 2019.

28. Vehtari A, Gelman A, Gabry J. Efficient implementation of leave-one-out cross-validation and WAIC for evaluating fitted Bayesian models. ArXiv Prepr ArXiv150704544. 2015.

29. Hawkins DM. The Problem of Overfitting. J Chem Inf Comput Sci. 2004;44: 1–12. doi:10.1021/ci0342472

30. Wilson RC, Collins AG. Ten simple rules for the computational modeling of behavioral data. eLife. 2019;8: e49547. doi:10.7554/eLife.49547

31. Korem N, Duek O, Ben-Zion Z, Kaczkurkin AN, Lissek S, Orederu T, et al. Emotional numbing in PTSD is associated with lower amygdala reactivity to pain. Neuropsychopharmacology. 2022; 1–9. doi:10.1038/s41386-022-01405-2

32. Korem N, Duek O, Spiller T, Ben-Zion Z, Levy I, Harpaz-Rotem I. Emotional State Transitions in Trauma-Exposed Individuals With and Without Posttraumatic Stress Disorder. JAMA Netw Open. 2024;7: e246813. doi:10.1001/jamanetworkopen.2024.6813

33. Staunton H, Willgoss T, Nelsen L, Burbridge C, Sully K, Rofail D, et al. An overview of using qualitative techniques to explore and define estimates of clinically important change on clinical outcome assessments. J Patient-Rep Outcomes. 2019;3: 16. doi:10.1186/s41687-019-0100-y

34. Higashi RT, Kruse G, Richards J, Sood A, Chen PM, Quirk L, et al. Harmonizing Qualitative Data Across Multiple Health Systems to Identify Quality Improvement Interventions: A Methodological Framework Using PROSPR II Cervical Research Center Data as Exemplar. Int J Qual Methods. 2023;22: 16094069231157345. doi:10.1177/16094069231157345

35. Kumar G, Basri S, Imam AA, Khowaja SA, Capretz LF, Balogun AO. Data Harmonization for Heterogeneous Datasets: A Systematic Literature Review. Appl Sci. 2021;11: 8275. doi:10.3390/app11178275

36. Levy D, Glimcher P. Comparing Apples and Oranges: Using Reward-Specific and Reward-General Subjective Value Representation in the Brain. J Neurosci. 2011;31: 14693–14707. doi:10.1523/JNEUROSCI.2218-11.2011

37. Han PKJ. Uncertainty and Ambiguity in Health Decisions. In: Diefenbach MA, Miller-Halegoua S, Bowen DJ, editors. Handbook of Health Decision Science. New York, NY: Springer; 2016. pp. 133–144. doi:10.1007/978-1-4939-3486-7_10

38. Reyna VF. A Theory of Medical Decision Making and Health: Fuzzy Trace Theory. Med Decis Making. 2008;28: 850–865. doi:10.1177/0272989X08327066

39. Attema AE, Bleichrodt H, L’Haridon O, Peretti-Watel P, Seror V. Discounting health and money: New evidence using a more robust method. J Risk Uncertain. 2018;56: 117–140. doi:10.1007/s11166-018-9279-1

40. Grubb MA, Li Y, Larisch R, Hartmann J, Gottlieb J, Levy I. The composition of the choice set modulates probability weighting in risky decisions. Cogn Affect Behav Neurosci. 2023;23: 666–677. doi:10.3758/s13415-023-01062-y

41. Gilaie-Dotan S, Tymula A, Cooper N, Kable JW, Glimcher PW, Levy I. Neuroanatomy Predicts Individual Risk Attitudes. J Neurosci. 2014;34: 12394–12401. doi:10.1523/JNEUROSCI.1600-14.2014

42. Grubb MA, Tymula A, Gilaie-Dotan S, Glimcher PW, Levy I. Neuroanatomy accounts for age-related changes in risk preferences. Nat Commun. 2016;7: 13822. doi:10.1038/ncomms13822

43. Schoemaker PJH. The Expected Utility Model: Its Variants, Purposes, Evidence and Limitations. J Econ Lit. 1982;20: 529–563.

44. Gong J, Zhang Y, Yang Z, Huang Y, Feng J, Zhang W. The framing effect in medical decision-making: a review of the literature. Psychol Health Med. 2013;18: 645–653. doi:10.1080/13548506.2013.766352

45. Kőszegi B, Rabin M. Reference-Dependent Risk Attitudes. Am Econ Rev. 2007;97: 1047– 1073. doi:10.1257/aer.97.4.1047

46. Lim RG. A Range-Frequency Explanation of Shifting Reference Points in Risky Decision Making. Organ Behav Hum Decis Process. 1995;63: 6–20. doi:10.1006/obhd.1995.1057

